# A versatile, high-efficiency platform for CRISPR-based gene activation

**DOI:** 10.1101/2022.07.21.501015

**Authors:** Amy J. Heidersbach, Kristel M. Dorighi, Javier A. Gomez, Ashley M. Jacobi, Benjamin Haley

## Abstract

CRISPR-mediated transcriptional activation (CRISPRa) is a powerful technology for inducing gene expression from endogenous loci with exciting applications in high throughput gain-of-function genomic screens and the engineering of cell-based models. However, current strategies for generating potent, stable, CRISPRa-competent cell-lines present limitations for the broad utility of this approach. Here, we provide a high-efficiency, self-selecting CRISPRa enrichment strategy, which combined with piggyBac transposon technology enables rapid production of CRISPRa-ready cell populations compatible with a variety of downstream assays. We complement this with a new, optimized guide RNA scaffold that significantly enhances CRISPRa functionality. Finally, we describe a novel, synthetic guide RNA tool set that enables transient, population-wide gene activation when used with the self-selecting CRISPRa system. Taken together, this versatile platform greatly enhances the potential for CRISPRa across a wide variety of cellular contexts.

## Main

Recent advances in genome engineering technology have enabled unprecedented opportunities for exploring the consequences of altered gene function or expression in a variety of model systems.^1,2^ Driving many of these efforts has been the adaptation of the microbial CRISPR/Cas9 system for use in eukaryotic organisms^3^. Cas9’s defining feature, as an easily programmable RNA-directed double stranded DNA (dsDNA) nuclease, has inspired the creation of genome-scale perturbation libraries and subsequent loss-of-function screens across hundreds of human cell lines^4–6^. These screens have proven invaluable for uncovering genotype and cell lineage-specific gene dependencies, which continue to inform basic as well as clinical research efforts^7^.

Cas9 can also be engineered for expanded use beyond the creation of targeted dsDNA breaks. The fusion of transcriptional repressor or activator domains to a nuclease-dead form of Cas9 (dCas9), enables CRISPR-mediated transcriptional interference (CRISPRi) or activation (CRISPRa), respectively^8–10^. CRISPRa is a compelling technology for the activation of endogenous gene expression in disease models or gain-of-function screens^11–13^. A host of activator domains and transgene expression systems have been engineered to enable the production of CRISPRa-competent cells^14^. However, current strategies for engineering CRISPRa transgenic cell-lines are inefficient, prone to silencing, and often necessitate a labor-intensive single-cell cloning process. Gene and cell line-dependent variability pose further limits on the scalability of CRISPRa.

Here, we provide a comprehensive platform based on the Synergistic Activation Mediator (SAM)^13^ CRISPRa concept, that takes advantage of a self-selection mechanism to create uniform, potent, and stable CRISPRa-competent cell populations without the need for clonal selection. In addition, we demonstrate the effectiveness of a new SAM-compatible single-guide RNA (sgRNA) variant that both improves the function of sub-optimal sgRNAs and enables activity from sgRNAs found to be inactive with earlier-generation scaffolds. We show that this new sgRNA format is not only capable of facilitating stable gene expression, but that it can also be used for transient target activation through a novel, chemically-synthesized guide RNA tool set. Altogether, this new, user-friendly platform maximizes the potential for CRISPRa across a breadth of cell-based contexts and genetic loci.

## Results

### A self-selecting CRISPRa strategy for the rapid generation of stable, high-efficiency CRISPRa cell populations

Several dCas9-activator concepts have been described^12^. In pilot experiments, we observed consistent evidence of target activation with the Synergistic Activation Mediator (SAM) system (data not shown), and selected this platform for optimization studies. The SAM system poses a challenge, however, owing to the size and number of discrete elements that must be introduced in order to create a stable CRISPRa-ready cell population. These include a dCas9-VP64 fusion protein, an MCP- (MS2 coat protein) p65-HSF1 co-activator fusion protein (MPH), and any number of selection markers. Combined, these components and their associated regulatory sequences exceed the conventional limit for efficient lentiviral packaging^15^, often necessitating a multi-vector delivery strategy^13,16^. The piggyBac transposon system^17^, on the other hand, allows for both a higher cargo capacity and the incorporation of multiple transgene cassettes within a single vector. PiggyBac-based strategies have been utilized for CRISPRa-based cell line generation^18,19^, but, similar to lentivirus, its use results in random genomic integration and functional heterogeneity within the cell population. The resulting low efficiency populations are often incompatible with demanding applications like functional genomic screening without the further derivation and characterization of high efficiency clones. Due to the laborious and time-consuming nature of this process we aimed to develop a simple and efficient bulk selection method to enrich for stable, uniform, and potent CRISPRa-expressing cell populations.

To this end, we designed a series of multi-component CRISPRa piggyBac vectors which employed individual selection strategies for the enrichment of transgenic cells (Fig. 1a). In each context, expression of the CRISPRa activator elements was driven by a human EF1*α* promoter, and this was complemented by a distinct mechanism for the transcription of a co-expressed puromycin resistance gene (puro^r^). Similar to previous studies, we created a *dual promoter* selection vector^19^ (Fig. 1a-top row) where puro^r^ was driven by an independent promoter (PGK) and a *single transcript* vector^20^ where puro^r^ was transcriptionally linked to the CRISPRa machinery (Fig. 1a-middle row). The theoretical selection pressure exerted by these strategies should be on maintaining transgene genomic integration in the case of the dual promoter vector and on sustained transgene expression in the case of the single transcript system (Fig. 1a-right column). As a readout for CRISPRa function we also incorporated a GFP reporter downstream of a self-activating (SA) promoter, which could be activated only in the presence of functional CRISPRa machinery and a co-expressed SA-targeting guide RNA. Building on the self-activating concept, we devised a third strategy, which we term *CRISPRa selection* (CRISPRa-sel), where the puro^r^ gene is driven by the self-activating promoter and linked to the GFP reporter (Fig. 1a-bottom row). Unlike the dual promoter or single transcript approaches, in the CRISPRa-sel context there is an absolute requirement for each cell to maintain functional CRISPRa in order to survive in the presence of puromycin.

**Fig. 1:**
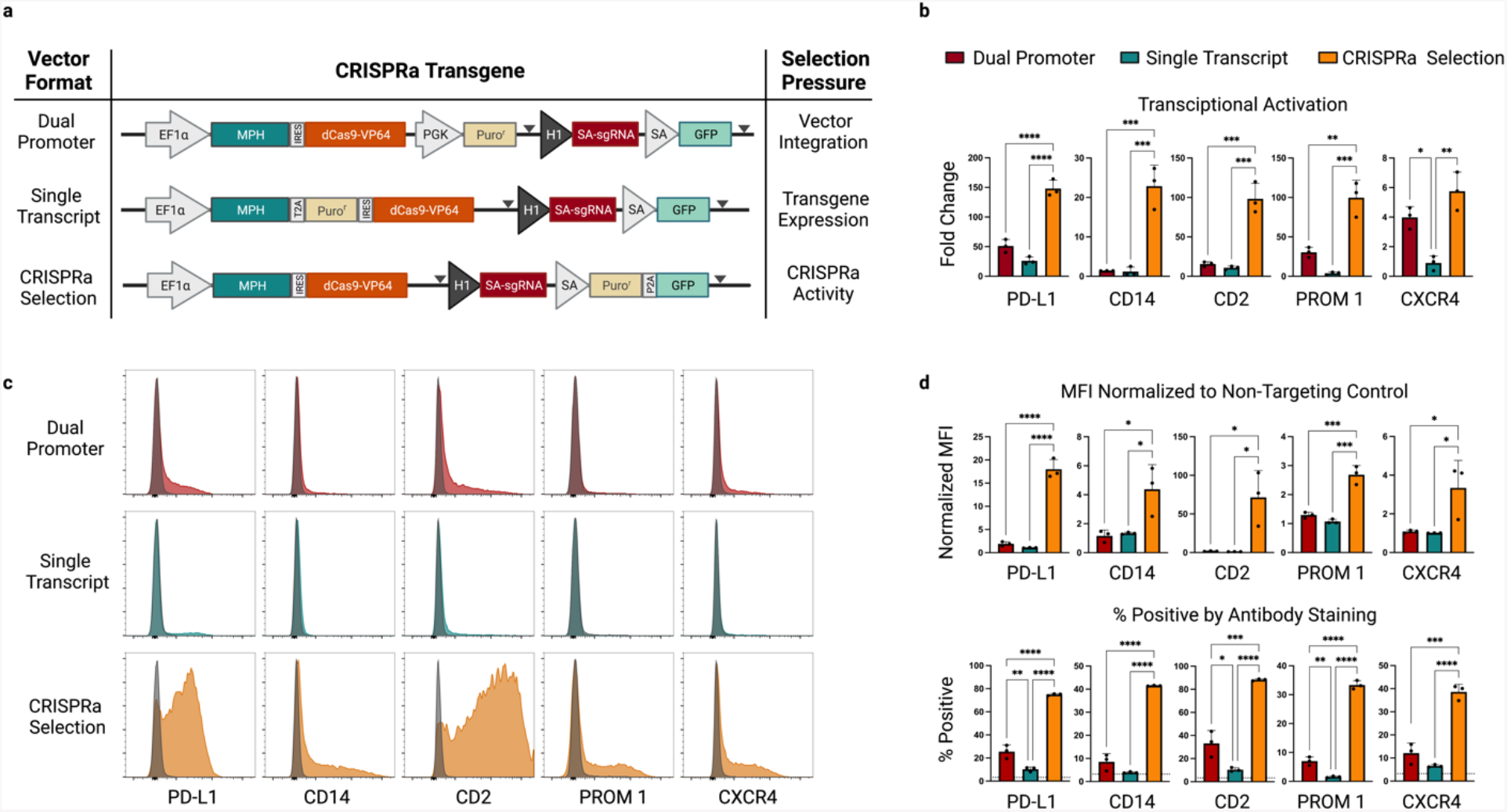
A self-selecting CRISPRa piggyBac vector for the rapid generation of stable, high-efficiency CRISPRa cell populations. **a**, Vector format and selective strategy for the evaluated piggyBac CRISPRa expression-reporter vectors. Expression of the MCP-P65-HSF1 (MPH) activator and dCas9-VP64 is driven by a constitutive, human EF1a promoter. A human H1 promoter drives constitutive expression of a sgRNA complementary to the self-activating (SA) promoter upstream of a GFP reporter. Expression of a puro^r^ gene is driven either by its own constitutive promoter (dual promoter), transcriptionally linked to the MPH/ dCas9-VP64 (single transcript) or under control of the CRISPRa dependent SA promoter (CRISPRa selection). Grey triangles indicate the location of LoxP sites. PiggyBac engineered K562 populations were generated in triplicate for each vector format and enriched with puro selection. sgRNAs complementary to the promoter proximal region of the indicated genes were cloned into a lentiviral vector context containing a mTagBFP2/zeocin selection cassette (Extended data 1a). Following transduction and zeocin selection target gene expression was evaluated by quantitative RT-PCR (qRT-PCR) **b**, and flow cytometry at day 14 post-infection (**c-d**). Representative histograms for each condition are overlaid with histograms from stained cell populations expressing a non-targeting control gRNA (**c**) (gray). Infections were performed in duplicate and averaged. (Median fluorescence intensity (MFI) was normalized to MFI of an antibody-stained sample expressing a non-targeting gRNA (**d, top**). Percentage of cells positive by antibody staining is presented (**d, bottom**) and background staining from a control sample expressing a non-targeting gRNA is indicated with a dashed horizontal line for each gene. Statistical comparison was performed by an unpaired 1-way ANOVA. * p<0.5, ** p<0.01, *** p<0.001. EF1a-Elongation factor alpha, GFP-green fluorescent protein, dCas9-vp64-nuclease dead spCas9+vp64 activator fusion, P2A-porcine teschovirus-1 2A self-cleaving peptide, HSF-heat shock factor, PD-L1-Programmed death-ligand1 (CD274), CD14-cluster of differentiation 14, CD2-Cluster of differentiation 2, Prom1-prominin-1 (CD133), CXCR4-C-X-C chemokine receptor type 4 (CD184).

We evaluated the relative efficiency of each selection strategy in the human K562 cell line. Following puromycin selection, the individual populations were infected with lentiviral vectors expressing SAM-compatible sgRNAs targeting the promoter proximal regions of five cell surface receptor genes (Extended Data Fig. 1a). Quantitative RT-PCR (qRT-PCR) data from each condition revealed consistently-improved gene activation with the CRISPRa-sel system relative to the other formats (Fig.1b). We further used flow cytometry to quantitatively assess cell surface protein expression on an individual cell level (Fig.1c,d; Extended Data Fig. 1b). This analysis further demonstrated a dramatic enhancement in gene activation with the CRISPRa-sel system both in terms of absolute protein expression, by way of normalized median fluorescence intensity (MFI), and percent positive-stained cells. While the dual promoter and single transcript systems showed highly heterogenous populations with only a small number of active cells, the CRISPRa-sel strategy resulted in a substantial improvement in the proportion of active cells, which in some cases achieved near population-wide activation (i.e. PD-L1 and CD2). To confirm the broad applicability of our findings, we expanded our analysis to two additional, unrelated human cell lines (Extended Data Fig. 1 c,d) where similar trends were observed. Importantly, potent endogenous gene activation with the CRISPRa-sel format suggested the self-activating circuit did not interfere with gene expression induced by separate, lentivirally-delivered sgRNAs.

In addition to endogenous target activation, we evaluated whether our integrated CRISPRa-dependent GFP reporter was effective at identifying CRISPRa-competent cells. To our surprise, we found that GFP intensity did not reliably correlate with endogenous target gene activation across the tested selection formats and cell lines (Extended Data Fig. 2). Although a trend towards correlation was observed for the dual promoter format (Extended Data Fig. 2a-left), there was high variability in the single transcript and CRISPRa-sel contexts (Extended Data Fig. 2a-center, right). To further evaluate the relationship between GFP expression and endogenous gene activation in the CRISPRa-sel context, we expanded our analysis to a second endogenous target gene (CD2) (Extended Data Fig. 2b) and observed similarly weak correlations. Despite this observation when analyzed in bulk, we wanted to determine if GFP expression could be used to facilitate the isolation of high-functioning single cell clones. We engineered the CRISPRa-sel system into four unrelated cell lines and following puro selection we sorted cell populations based on high, medium, or low GFP expression via fluorescence activated cell sorting (FACS) (Extended Data Fig. 2c-d). From these sorted populations, single-cell clonal lines were derived, and upon expansion were transduced with sgRNAs targeting distinct endogenous genes (PD-L1, CXCR4) or a non-targeting control. Interestingly, while relative GFP expression levels were maintained in the clones post-expansion (Extended Data Fig. 2c-left), there was no clear relationship between reporter expression and endogenous target activation in three of four cell lines evaluated (Extended Data Fig. 2c-center/right). These data suggest that, in the context of the CRISPRa-sel system, selection with a CRISPRa dependent fluorescent reporter is not a broadly applicable strategy for further enrichment of CRISPRa-competent populations, beyond what is achieved with puromycin selection. While other groups have reported successful enrichment with fluorescent CRISPRa responsive reporters^20^, our data suggest such strategies are potentially more useful in the context of low efficiency systems like the dual promoter or single transcript formats where the number of active cells in the population is low and the functional difference between active and inactive cells is high. On the other hand, GFP-based reporters may not be sensitive enough to discriminate effectively between cells within more uniform CRISPRa-sel derived cell populations. Therefore, we focused on antibiotic-selected populations for the remainder of our platform optimization efforts.

### SAM guide RNA scaffold optimization for enhanced CRISPRa activity

Subtle changes in scaffold sequence and structure have been shown to affect guide RNA function^13,16,21,22^ and we reasoned that the conventional SAM-2.0 scaffold could be re-engineered to improve activity. The MPH activator utilized by the SAM system binds to two separate MS2 aptamers within the SAM-2.0 sgRNA; one in the tetraloop and one in stem loop two (Extended Data Fig. 3a). Focusing on the tetraloop, we used rational design to create several new SAM-compatible scaffold variants (Extended Data Fig. 3a-b, Fig. 2a). Previous reports have indicated that Pol-III-based guide expression can be enhanced by removing a poly U tract in the tetraloop, which can serve as a premature transcriptional termination sequence.^21,22^ (Extended Data Fig. 3b [GNE-1]). Additionally, we hypothesized that increasing the stability or accessibility of the MS2 aptamer segment within the tetraloop could encourage greater associations with MPH complexes, further improving CRISPRa efficiency. To explore these possibilities, we coupled poly U deletion with an alternate, GC-rich stem extension sequence proximal to the MS2 aptamer (Extended Data Fig. 3 [GNE-2])^21^. Finally, we combined both stem extension features with the removal of a bulge sequence directly adjacent to the MS2 aptamer (Extended Data Fig. 3 [GNE-3]).

**Fig. 2:**
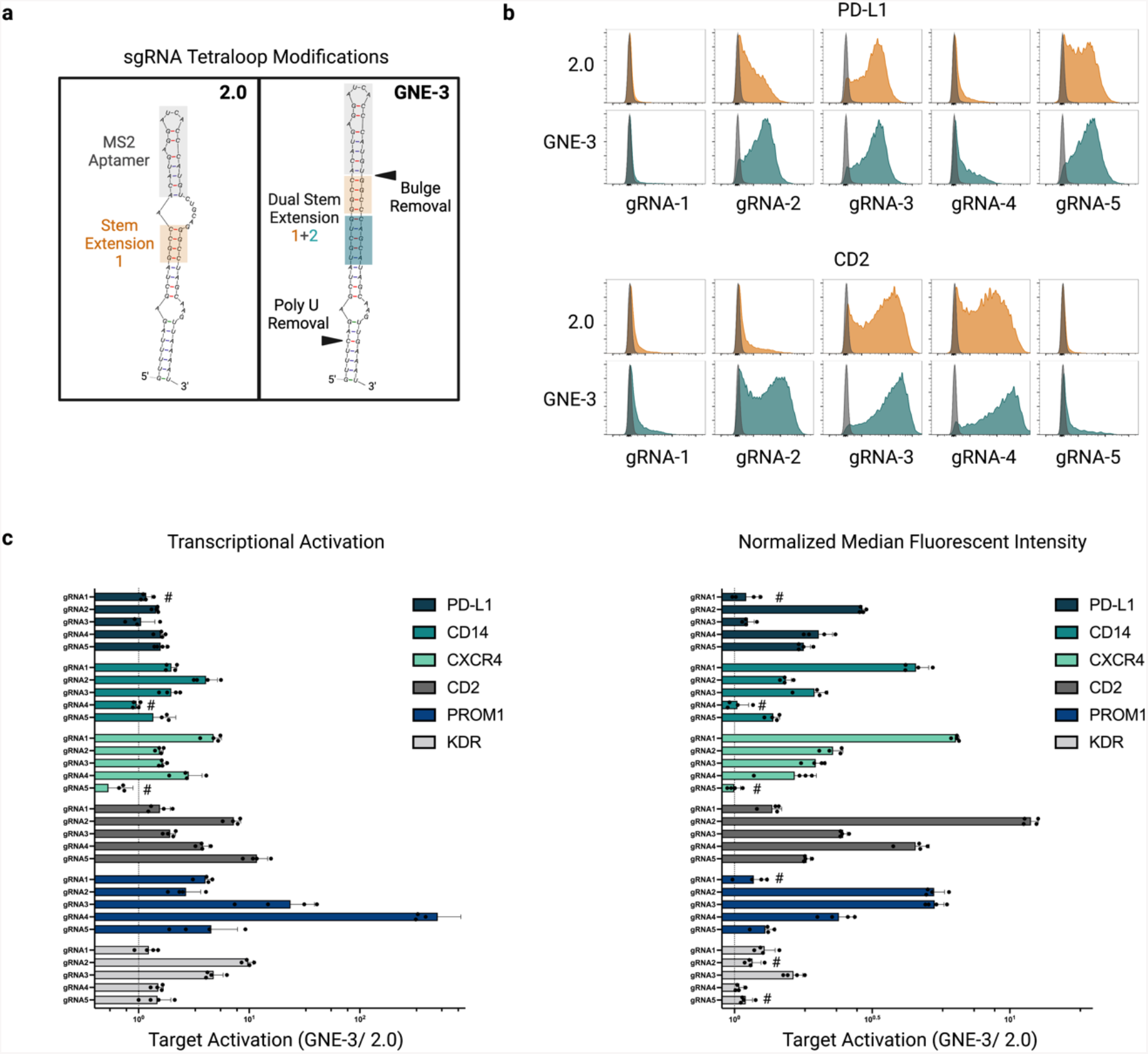
Relative CRISPR activation efficiency of sgRNAs containing an optimized MS2 aptamer containing scaffold. **a**, Structure diagram of the MS2-aptamer containing tetraloop in the 2.0 sgRNA format^13^ (left) or an optimized tetraloop structure (right and Extended Data Fig. 3). The optimized GNE-3 tetraloop contains an additional stem extension and removal of a polyU tract^21^. Additionally, the bulge region connecting the MS2 aptamer and stem extension region 1 in the 2.0 format has been removed. **b**, Flow cytometric analysis of target gene activation by sgRNAs with either a 2.0 (orange) or GNE-3 (teal) scaffold context. Representative histograms of analyzed K562 CRISPRa-sel populations infected with 5 distinct spacer sequences targeting the promoter proximal region of PD-L1 (top) or CD2 (bottom). Populations infected with a non-targeting sgRNA sequence overlaid (gray). **c**, Activation of 6 target genes by GNE-3 sgRNAs normalized to the activation efficiency of the same spacer sequence in a 2.0 format (dashed line). Normalized gene activation was evaluated in zeocin selected populations by qRT-PCR (left) at day 14 post-sgRNA infection or by flow cytometry (right) at day 10 post-sgRNA infection. n=4 replicates per sgRNA.

To evaluate the relative efficiency of these scaffolds, we lentivirally-transduced CRISPRa-sel engineered K562 populations with sgRNAs targeting three endogenous genes (PD-L1, CD14, or KDR) in either the SAM-2.0 scaffold format or one of our three novel variants (Extended Data Fig. 3c). By flow cytometry, higher target expression was observed with several of the new scaffold variants, but GNE-3 showed the most consistent improvement over 2.0, both in terms of gene product levels (normalized MFI) and the percentage of activated cells across the population. We subsequently expanded our comparison of the 2.0 and GNE-3 scaffolds to include six cell surface receptor genes, using five unique sgRNAs per gene, to account for gene and spacer-specific variability. Analysis of target transcript (qRT-PCR) and protein (flow cytometry) expression (Fig. 2b-c; Extended Data Fig. 4a-b) revealed a broad enhancement of target activation with the GNE-3 scaffold versus the 2.0 backbone, with several sequences achieving between 5-10-fold improved gene induction with the GNE-3 variant. To confirm that the GNE-3 scaffold was beneficial in other cell contexts, we expanded our analysis to two additional cell lines. As before, we found activation of PD-L1, as measured by cell surface staining in 293T and Jurkat cells, (Extended Data Fig. 4c) was consistently higher with the GNE-3 scaffold. Taken together these data suggest that the GNE-3 scaffold improves both the breadth and magnitude of gene activation across a variety of spacer, target and cellular contexts. In addition, we found that relative target gene activation was largely consistent when comparing transcript level or cell surface protein stain (Extended Data Fig. 4d) for most targets, and therefore chose to move forward with validated flow cytometry assays for subsequent experiments owing to the quantitative nature of this assay at both the population and individual cell levels.

### CRISPRa-sel promoter optimization and evaluation in a panel of human cell lines

While the combination of our CRISPRa-sel system with the GNE-3 scaffold demonstrated improvement in overall CRISPRa efficiency, we continued to observe variable target activation across cell lines (Fig. 3a-c [EF1*α*], Extended Data Fig. 5 [EF1*α*]). The strength of Pol-II promoters, which drive expression of the CRISPRa machinery, can differ dramatically across cell types^23^ potentially contributing to the context-dependent efficacy of CRISPRa (Fig. 3a). To evaluate how promoter use impacts the efficiency of the CRISPRa-sel system, we engineered a panel of three cell lines (K562, 293T and Jurkat) with the original EF1*α*-based CRISPRa-sel vector or versions that incorporated three distinct cytomegalovirus (CMV)-derived Pol-II promoter variants (CBh, CMV, and CAG) (Fig. 3a-c.; Extended data 5) to drive expression of the activator machinery.

**Fig. 3:**
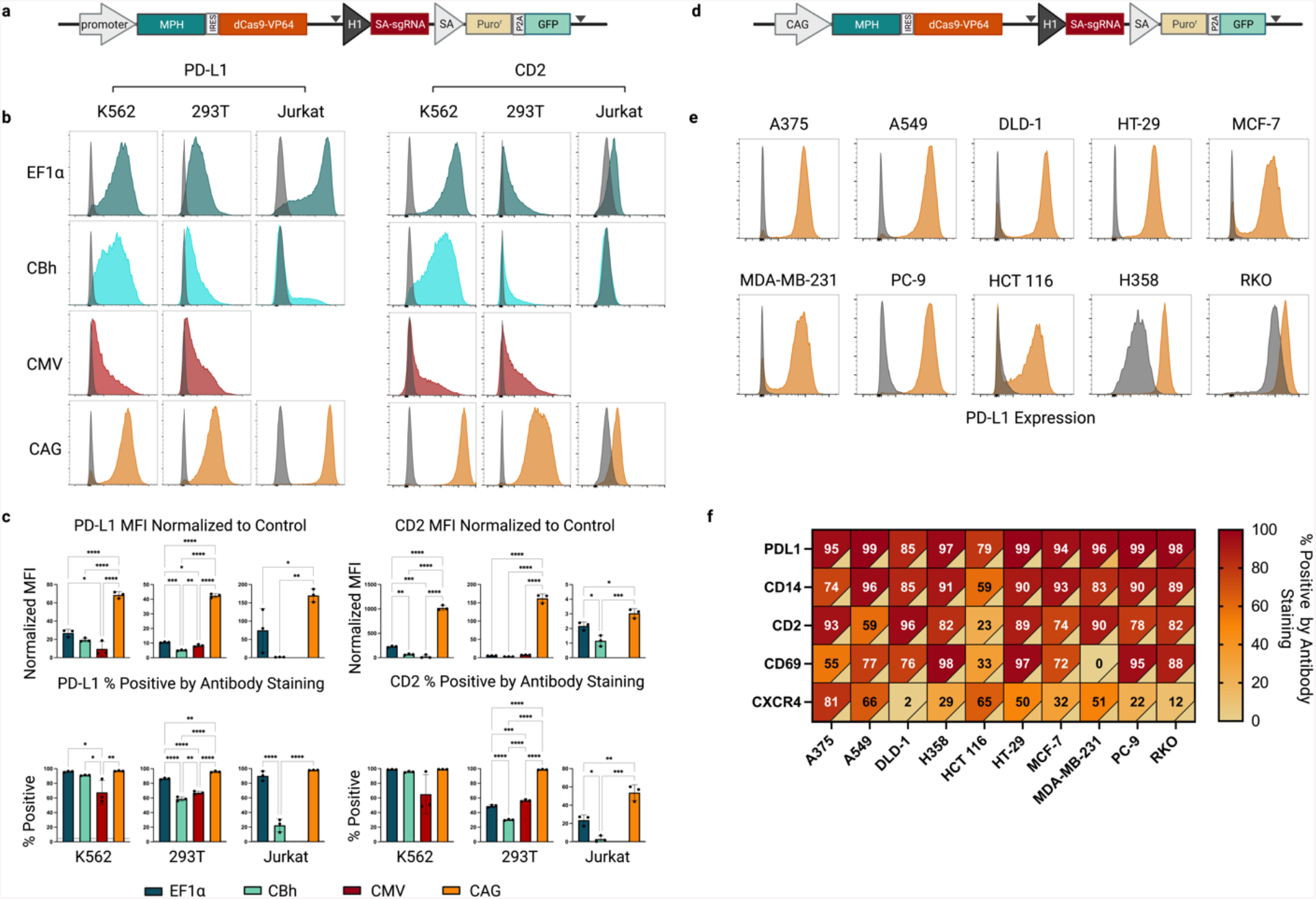
Promoter optimization and application of the CRISPRa-sel strategy across a panel of commonly used cell lines. **a**, Schematic representation of the CRISPRa-sel vector indicating the location of the promoter driving expression of the MPH/dCas9-VP64 transcript. **b-c**, Activation of PD-L1 (left) or CD2 (right) target genes evaluated by flow cytometry in K562, 293T and Jurkat cell lines engineered with CRIPSRa-sel piggyBac vectors utilizing an EF1a (teal), CBh (aqua), CMV (maroon) or CAG (orange) promoter 14 days post-infection with a GNE-3 sgRNA. **(b)** Activation displayed by representative histograms overlaid with expression profiles from cells infected with a non-targeting sgRNA (gray). **(c)** Normalized median fluorescence intensity (MFI) (top) or percentage positive (bottom) of indicated genes/cell populations by antibody staining. The percent positive of stained control populations infected with a non-targeting sgRNA are indicated by a dashed horizontal line. (Note: *CMV CRISPRa-sel Jurkat populations did not grow out efficiently and were not included in the analysis*.) **(d)** Schematic representation of the CAG CRISPRa-sel piggyBac vector. **(e)** Representative flow cytometric histograms of PD-L1 activation across 10 commonly used cell lines engineered with a CAG-driven CRISPRa-sel piggyBac vector and PD-L1 targeting GNE-3 sgRNA. **(f)** Heatmap representing the percent positive of 5 target genes (PD-L1, CD14, CD2, CD69 and CXCR4) across 10 CAG CRISPRa-sel engineered cell lines (A375, A549, DLD-1, H358, HCT 116, HT-29, MCF-7, MDA-MB-231, PC-9 or RKO). Percent positive of stained cell populations expressing a non-targeting sgRNA represented colorimetrically in the lower right corner of each cell. Gene-activating or control guides were expressed using dual sgRNA lentivectors (supplemental methods). CRISPRa cell populations generated in triplicate and infected with indicated sgRNAs in technical duplicates which were averaged before statistical comparison was performed by an unpaired 1-way ANOVA. * p<0.5, ** p<0.01, *** p<0.001. Grey triangles indicate the location of LoxP sites. CBh -Chicken β-actin hybrid promoter, CMV- human cytomegalovirus immediate-early gene enhancer/promoter or CAG- ***C***MV enhancer-chicken β-***a***ctin-rabbit β-***g***lobin synthetic hybrid promoter.

Attempts to engineer CRISPRa-sel populations were successful in all but one cell line context (Jurkat + CMV-CRISPRa-sel) (Fig. 3b), in which only a low number of slow growing clones were recovered following puromycin selection. To evaluate the relative efficacy of each promoter, populations were transduced with GNE-3 sgRNAs targeting PD-L1 or CD2. Unlike the more heterogeneous activation observed with the EF1*α*, CBh, and CMV promoters, the CAG promoter induced distinctly uniform and potent gene expression for each of the tested cell lines and targets (Fig. 3b-c). We expanded our assessment to include three additional endogenous targets (CD14, CXCR4 and CD69) and saw comparable results (Extended Data 5a-b). Importantly, this demonstrated that population-wide CRISPRa was achievable with limited cell culture manipulation steps beyond bulk antibiotic selection.

In order to confirm the broad utility of the CAG-CRISPRa-sel and GNE-3-sgRNA system, we engineered an additional panel of ten commonly used cell lines (Fig. 3d-f). After bulk selection of the CAG-CRISPRa-sel transgenic cell lines, introduction of a PD-L1-specific sgRNA led to strong, uniform target induction (∼79-99% of the cell population) (Fig. 3e-f). We then expanded this analysis to four additional target genes per cell line, and while we observed some context-dependent variability for individual genes, robust activation in ≥75% of the cell population was seen in the majority of conditions. Notably, beyond activating genes with little or no background expression, we were able to induce population-wide upregulation of genes with high basal expression (Fig. 3e [H358^24^],[RKO^25^]). Taken together these data indicate that the CAG-CRISPRa-sel system in conjunction with the GNE-3 scaffold greatly enables the utility of stable CRISPRa across a breadth of cell backgrounds and target genes.

### Optimized, multi-format synthetic guide RNAs for transient CRISPRa

Synthetic guide RNAs can be generated quickly and have proven effective for Cas9-mediated gene disruption purposes ranging from the creation of *in vitro* and *in vivo* models to arrayed genetic screens^26^. While synthetic gRNAs have previously been applied in the context of CRISPRa^27^ thus far they have not been widely adopted possibly due to their low efficiency with sub-optimal CRISPRa systems. The production of synthetic, high-efficiency, SAM-compatible guides has presented technical challenges. Until recently, dual MS2 aptamer-containing sgRNAs, like the ∼160 nucleotide GNE-3 spacer sequence and scaffold, exceeded the length of reliable direct synthesis methodology ^28^. As an alternative approach, the use of easier-to-synthesize two-part gRNAs (crRNA + tracrRNA scaffold) is an attractive possibility. The design of these guides, however, must allow for efficient strand annealing while maintaining the structure of the MS2 aptamer loops^26^. In addition, any synthetic guide RNA, regardless of format, needs to be stable enough throughout the delivery, dCas9 association, and target binding processes to induce measurable gene activation. Given recent advances in RNA synthesis and chemical stabilization, and to yet further expand the utility of CRISPRa, we set out to develop an optimized GNE-3-based synthetic gRNA platform.

To evaluate the impact of chemical modifications on the efficiency of CRISPRa induced by transient delivery of synthetic guides in cultured cells, we synthesized a set of sgRNAs based on the GNE-3 scaffold targeting four endogenous genes (PD-L1, CD14, CD2, CXCR4) with or without modified stabilizing nucleotides^29^. Individual unmodified sgRNAs were compared to identical sgRNAs containing three terminal phosphorothioated 2’ O-methyl ribonucleotides at both the 5’ and 3’ ends (Extended Data Fig. 6a). Three days after electroporation into a CAG-CRISPRa-sel-engineered K562 population, we observed clear evidence of gene activation. We found that the modified sgRNAs demonstrated a clear advantage over the unmodified guides across all targets evaluated (Extended Data Fig. 6b-c). To our surprise, activation with the transient modified synthetic sgRNAs was qualitatively similar in some cases to stable sgRNA expression, with near-population-wide expression achieved for two of four target genes.

We next sought to determine if the GNE-3 sgRNA variant also outperformed the 2.0 scaffold in a synthetic context. To this end we generated identical end-modified sgRNAs for the 2.0 variant. Direct comparison in the CAG-CRISPR-sel K562 model demonstrated a general trend towards higher activation with the GNE-3 sgRNAs, although the differential was somewhat reduced compared to the stable sgRNA context (Extended Data Fig. 7).

User accessibility of synthetic guide RNA-mediated CRISPRa could be enhanced by lowering the cost and technical skill required for reagent synthesis. In principle, this could be achieved by minimizing the length of the guide RNA segments with a more native, annealed two-part crRNA-tracRNA format. In order to create synthetic material that permitted crRNA and tracrRNA hybridization while maintaining the GNE-3 scaffold loop structure, we developed two distinct concepts (Extended Data Fig. 8a). In format 1, strand 1 includes the spacer sequence and a segment of the GNE-3 MS2 containing tetraloop (Extended Data Fig. 8a-teal), which anneals to strand 2 containing the final portion of the tetraloop as well as stemloop 1, stemloop 2 (with the second MS2 aptamer) and stemloop 3. Separately, in format 2, strand 1 exclusively comprises the spacer plus a short region (Extended Data Fig. 8a-orange) with complementarity to strand 2. Strand 2 of this format encodes the majority of the tetraloop and stemloops 1-3. All RNA oligonucleotides contain 5’ and 3’ stabilizing modifications similar to our optimized synthetic GNE-3 sgRNA. We incorporated identical spacer sequences within both formats and evaluated their relative effectiveness for activating four separate genes within CAG-CRISPRa-sel K562 cells. When we analyzed target activation by flow cytometry three days post-electroporation we saw higher gene activation with format 1 across all targets (Extended Data Fig. 8b-c), and this format became a focus for follow-up studies.

Recently, a two-part, SAM-compatible guide RNA system has been described and made commercially available^27^. Unlike the GNE-3 guide RNAs described herein, the commercial product contains fewer phosphorothioated 2’ O-methyl ribonucleotides and has only a single MS2-modified element within stemloop 2, the MS2 sequence within the tetraloop being notably absent (Fig. 4a-top). In order to evaluate the relative functionality of these synthetic guide RNAs, we compared GNE-3 sgRNAs and format 1 two-part guide RNAs to the commercially available synthetic guide RNA format (1X MS2 two-part) in CAG-CRISPRa sel-engineered K562 and 293T populations (Fig. 4 and Extended Data Fig. 9). We found that the GNE-3 sgRNA and two-part formats generally outperformed the single MS2 containing guide (Fig. 4 and Extended Data Fig. 9) with the GNE-3 sgRNA format providing the most consistent and potent activation across all tested contexts. The differential across guides was particularly pronounced in lower activity conditions (Fig 4b-c, Extended Data Fig. 9a-b gRNA-1). Only under circumstances of high CRISPRa activity, such as in 293T cells, could measurable induction be achieved with all of the evaluated 2-part and sgRNA variants (Fig 4d-e, gRNA-3/gRNA-4, Extended Fig 9).

**Fig. 4:**
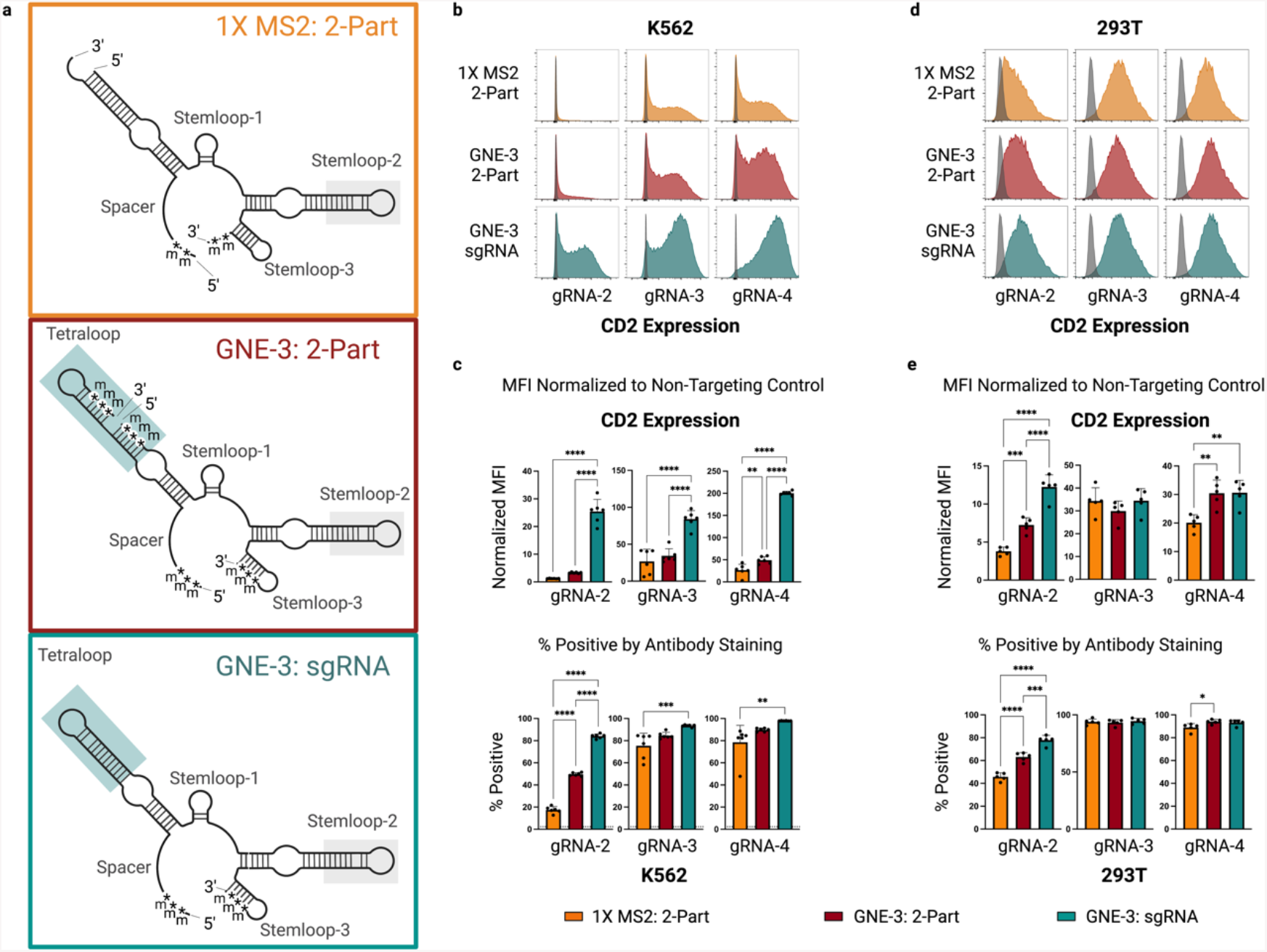
Evaluation of CRISPRa synthetic guide formats across 2 cell lines. **a**, Structure diagrams of a commercially available, chemically modified, 2-part synthetic gRNA containing a single MS2 aptamer loop (top-orange); a modified, 2-part format 1 synthetic gRNA containing a GNE-3 scaffold (center-maroon); or a modified sgRNA with a GNE-3 scaffold (bottom-teal). Blue boxes highlight the MS2 apatmer-containing GNE-3 tetraloop and grey boxes indicate the MS2-apatmer on stemloop 2. **b-e** CRISPR-mediated transcriptional activation of a CD2 target gene in two CAG-CRISPRa-sel engineered cell lines (K562 or 293T) by electroporated modified, synthetic gRNAs in the formats depicted in **(a)**. CD2 target expression by 3 spacer sequences in an engineered K562 cell line assessed by flow cytometry. CD2 expression displayed by representative histograms overlaid with a control population **(b)** or summarized by median fluorescent intensity normalized to a non-targeting control **(c-upper)** or percent positive **(c-lower)**. Percent positive of a stained control population infected with a non-targeting sgRNA are indicated by a dashed horizontal line. **d-e**, CD2 target activation by synthetic gRNA formats as in b-c but in a 293T cell line. Flow cytometry performed 3 days after synthetic guide delivery. Statistical comparison between guide formats was performed by an unpaired 1-way ANOVA. * p<0.5, ** p<0.01, *** p<0.001. n=6 for K562, n=5 for 293T. m=2’-O methyl. *= phosphorothioate linker.

## Discussion

The potential for any genome engineering technology is limited by the breadth of cell types and loci for which it can be applied. By incorporating a unique self-selecting transgenic approach with enhanced SAM-compatible guide RNA scaffolds, we have demonstrated that robust, population-wide CRISPRa is achievable across a diverse panel of target genes and cell lines, all with minimal cell manipulation steps. In addition, we show that synthetic guide RNAs can be employed for highly-efficient, short term gene activation, in some cases with population-wide efficacy. While this platform is expected to be broadly applicable, the required plasmid transfection process may limit use in cell types that are sensitive to foreign DNA or difficult to transfect with large plasmids. Adaptation of the self-selection concept with viral vector-based delivery could circumvent this bottleneck.

Advancements in gene activator technologies are inevitable. With this in mind, we anticipate that self-selecting circuits will be compatible with future transcriptional and epigenetic modifier fusion proteins or extended Cas family member usage^1^. This will be critical for expanding the target space available for CRISPRa and for potentially enhancing gene expression at loci that show weak or modest induction with the SAM activator machinery.

## Methods

### Cell culture, electroporation, transfection

Cell line specific culture and manipulation protocols described in supplemental methods. All parental cell lines were sourced from the Genentech cell bank (gCell) where they were maintained under mycoplasma free conditions and authenticated by STR profiling. FACS sorting and subsequent clonal derivation/analysis presented in extended data 2 c,d was performed by WuXi AppTech.

### Lentiviral production/transduction

sgRNA expressing and lentiviral packaging plasmids (VSVg/Delta8.9) were transiently co-transfected into 293T cells with Lipofectamine 2000. Lentiviral supernatants were harvested at 72 hours and filtered through a 0.45 µm PES syringe filter (Millipore). Transduction with lentivirally encoded guide RNAs performed as described in supplemental methods with cell line specific protocols. 3 days following lentiviral infection, cells were started on zeocin selection at cell line specific concentrations (supplemental methods) in order to select for guide RNA expressing cells. Prior to gene expression analysis, uniform selection of gRNA infected populations was confirmed by flow cytometric analysis of the co-expressed mTagBFP2 reporter.

### Flow cytometry

Antibody staining performed using manufacturers recommended protocols and described in supplemental methods. Data collection performed on BD FACS Celesta or BD FACS Symphony machines and analyzed by FlowJo 2 10.8.0. Gating strategy indicated in Extended Data Fig. 1b. Live cell populations were gated using FSC and SSC profiles. Where relevant, lentivirally transduced cells specifically were examined by gating on mTagBFP2 positive populations. If cell populations were selected to greater than >95% mTagBFP2 positive then this gating step was omitted for some analyses. Populations were defined by gates established as indicated with 2 parameter pseudocolor plots (Extended Data Fig. 1b) with identical control cell lines expressing a non-targeting control guide RNA and stained/collected in parallel.

### qRT-PCR

RNA extraction performed with a Quick-RNA 96 well kit (Zymo). cDNA generation performed with a high-capacity cDNA synthesis kit using random primers and RNase inhibitor (Thermo) following recommended protocols. Quantitative RT-PCR performed with an ABI QuantStudio 7 Flex real time PCR system. Relative quantification/fold change (2^-ΔΔCT) analysis was performed by QuantStudio software. A GAPDH control gene used for normalization purposes.

### Synthetic gRNA electroporation/transfection

Direct synthesis and QC of the novel modified sgRNA and 2-part guide RNAs was performed by IDT (https://www.idtdna.com/pages). All synthetic gRNAs were resuspended in Nuclease-Free Duplex Buffer (30 mM HEPES, pH 7.5; 100 mM potassium acetate) (IDT). Commercially available modified, synthetic 2-part guide RNAs containing a single MS2 aptamer loop purchased from Horizon inc. (https://horizondiscovery.com/).

2-part crRNA and tracrRNA oligonucleotides were combined at equimolar ratios prior to a denaturation/annealing protocol (95°C 5”; cool to room temp 2°/sec). sgRNAs were also treated by heat denaturation prior to use. Cell line specific synthetic guide delivery protocols detailed in supplemental methods.

### Data/Statical analysis

Statistical tests performed as indicated in figure legends for each experiment. Error bars represent standard deviation from the mean. Data was analyzed using PRISM and/or excel software. Bar plots/scatter plots and heatmaps were generated using PRISM.

### RNA structure prediction

RNA folding performed using mFold^30^ or bifold (https://rna.urmc.rochester.edu/RNAstructureWeb/Servers/bifold/bifold.html) algorithms.

### Figure production

Figure elements produced in Excel (Microsoft), Flowjo (Becton Dickson) and PRISM (Graphpad Software). Final figures created with BioRender.com.

## Supporting information

Supplemental Methods (Antibodies, qPCR probes, Full vector sequences, gRNA sequences and cell line information)

## Acknowledgements

We would like to acknowledge JP Fortin, Colin Watanabe, Søren Warming, Anqi Zhu, Sandra Melo, Yassan Abdolazimi, Nadia Martinez-Martin, Clark Ho, Fabiola Juárez and Letty Marroquin for thoughtful discussions and manuscript support.

## Disclaimers

A.H., K.D., and B.H. are full time employees of Genentech, Inc. and shareholders of Roche. Products and tools supplied by IDT are for research use only and not intended for diagnostic or therapeutic purposes. Purchaser and/or user is solely responsible for all decisions regarding the use of these products and any associated regulatory or legal obligations. J.A.G and A.M.J are employees of Integrated DNA Technologies, which offers reagents for sale similar to some of the compounds described in the manuscript.

## Data and Materials Availability Statement

The datasets generated during and/or analyzed during the current study are available from the corresponding author on reasonable request. Biological materials will be provided to requesters through a material transfer agreement. Vector and guide RNA sequences are provided in supplemental methods. Synthetic guide RNAs can be purchased through IDT.

## Figures and Figure Legends

**Extended Data Fig. 1:**
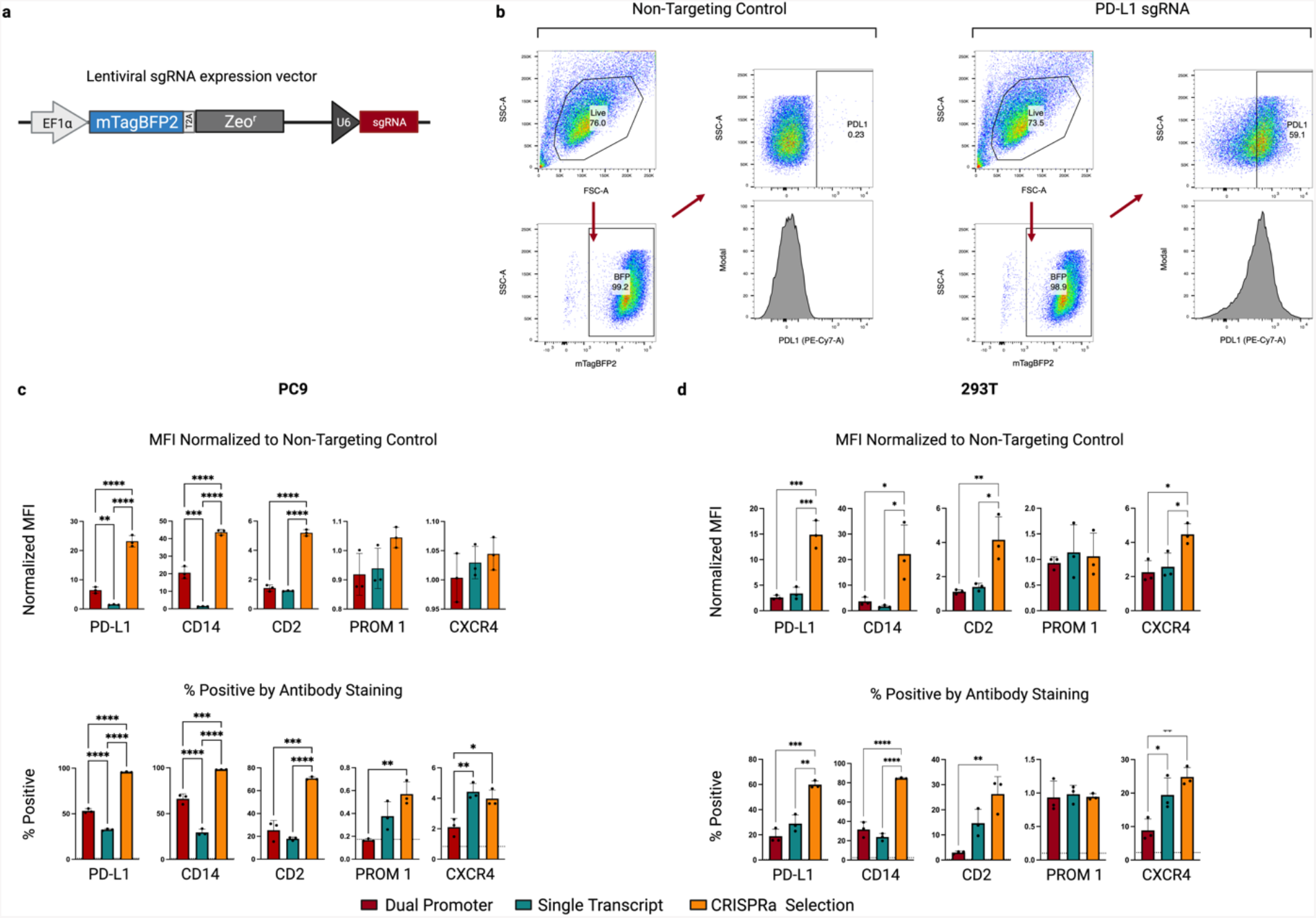
Comparison of CRISPRa piggyBac vector systems in additional cell lines. **a**, Schematic representation of the lentiviral sgRNA expression vector containing a mTagBFP2 marker and zeocin resistance (Zeo^r^) selection cassette. **b**, Representative gating strategy for flow cytometric analysis shown in CRISPRa-sel populations expressing either a non-targeting control (left) or PD-L1 targeting sgRNA (right). Live populations were identified as indicated based on SSC-A (side scatter) and FSC-A (forward scatter) profiles and sgRNA expressing cells were identified by expression of the mTagBFP2 florescent protein. Positive population gates were defined in a control sample stained in parallel. **c**, Flow cytometric analysis of CRISPRa mediated gene expression in cell populations generated with three CRISPRa piggyBac systems utilizing distinct selection strategies (Fig.1). Analysis performed 14-25 days post lentiviral transduction of sgRNAs complementary to the promoter proximal regions of the indicated genes (PD-L1, CD14, CD2, PROM1 or CXCR4) for PC-9, or **d**, 293T cells. (Median fluorescence intensity (MFI) was normalized to MFI of an antibody-stained sample expressing a non-targeting gRNA (top). Percent antibody positive is presented (bottom) and background staining from a control sample expressing a non-targeting gRNA is indicated with a dashed horizontal line for each gene. Cell populations generated in triplicate. sgRNAs infected in duplicate and averaged prior to statistical comparison with an unpaired 1-way ANOVA. * p<0.5, ** p<0.01, *** p<0.001.

**Extended Data Fig. 2:**
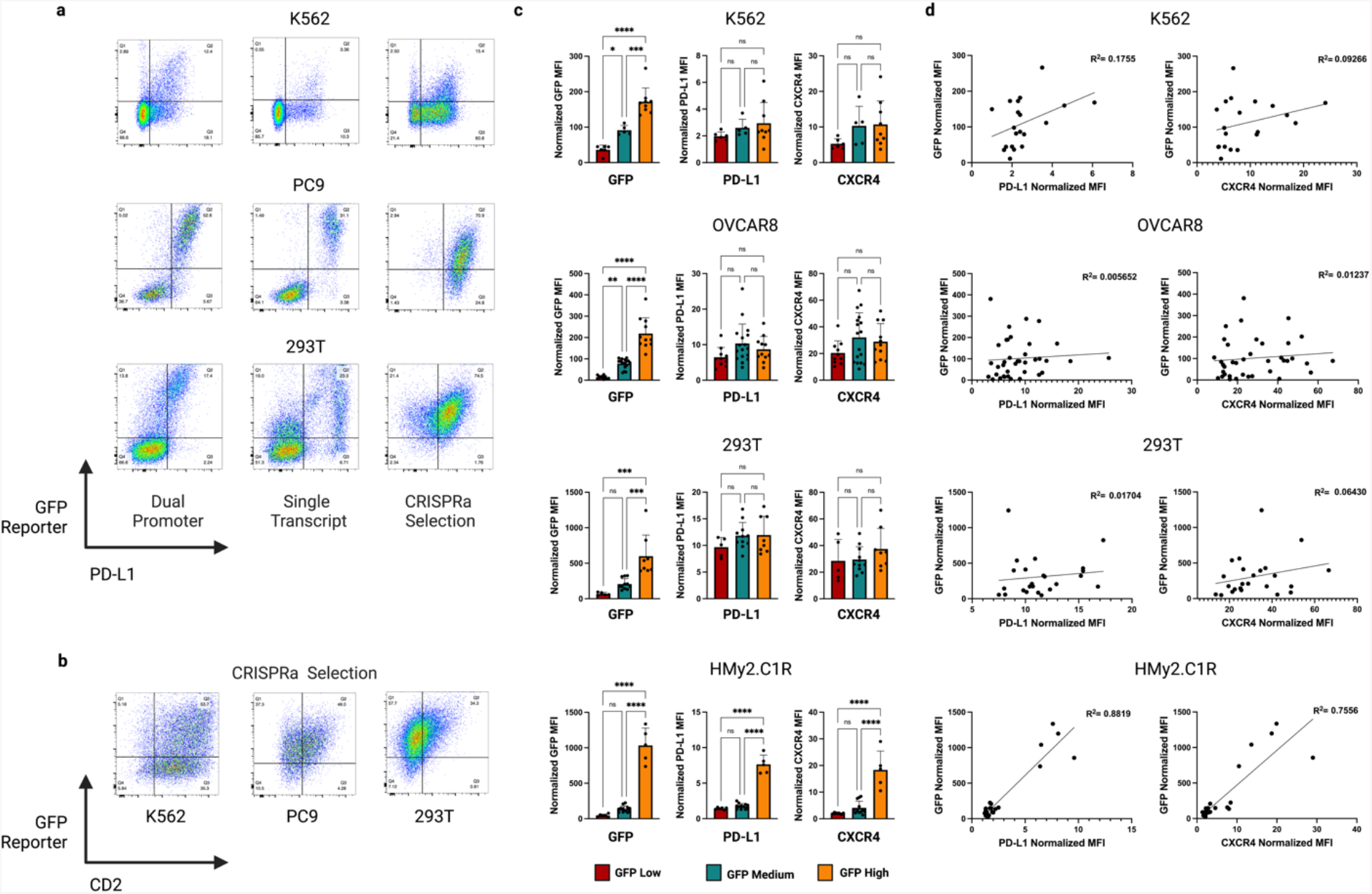
An integrated CRISPRa dependent GFP reporter is an inconsistent marker of CRISPRa efficiency across multiple cell lines. **a**, Flow cytometric analysis of GFP CRISPRa reporter vs endogenous PD-L1 target gene activation across three CRISPRa piggyBac formats in three cell lines (Fig. 1). **b**, GFP CRISPRa reporter vs endogenous CD2 activation in three cell lines engineered with a CRISPRa-sel piggyBac. **c**, Flow cytometric analysis of GFP CRISPRa reporter vs two endogenous CRISPRa target genes (PD-L1, CXCR4) in clones derived from CRISPRa-sel populations pre-sorted on GFP expression using flow assisted cell sorting (FACS) in four cell lines. Bar graphs of GFP median fluorescence intensity (MFI) in clones normalized to parental cell line (left). Target gene expression in engineered clones infected with an endogenous gene targeting sgRNAs (PD-L1 or CXCR4) and normalized to non-targeting control gRNA (middle/right). **d**, Scatter plots showing correlation of normalized MFI for CRISPRa dependent GFP reporter vs endogenous target gene activation. R squared for simple linear regression analysis indicated.

**Extended Data Fig. 3:**
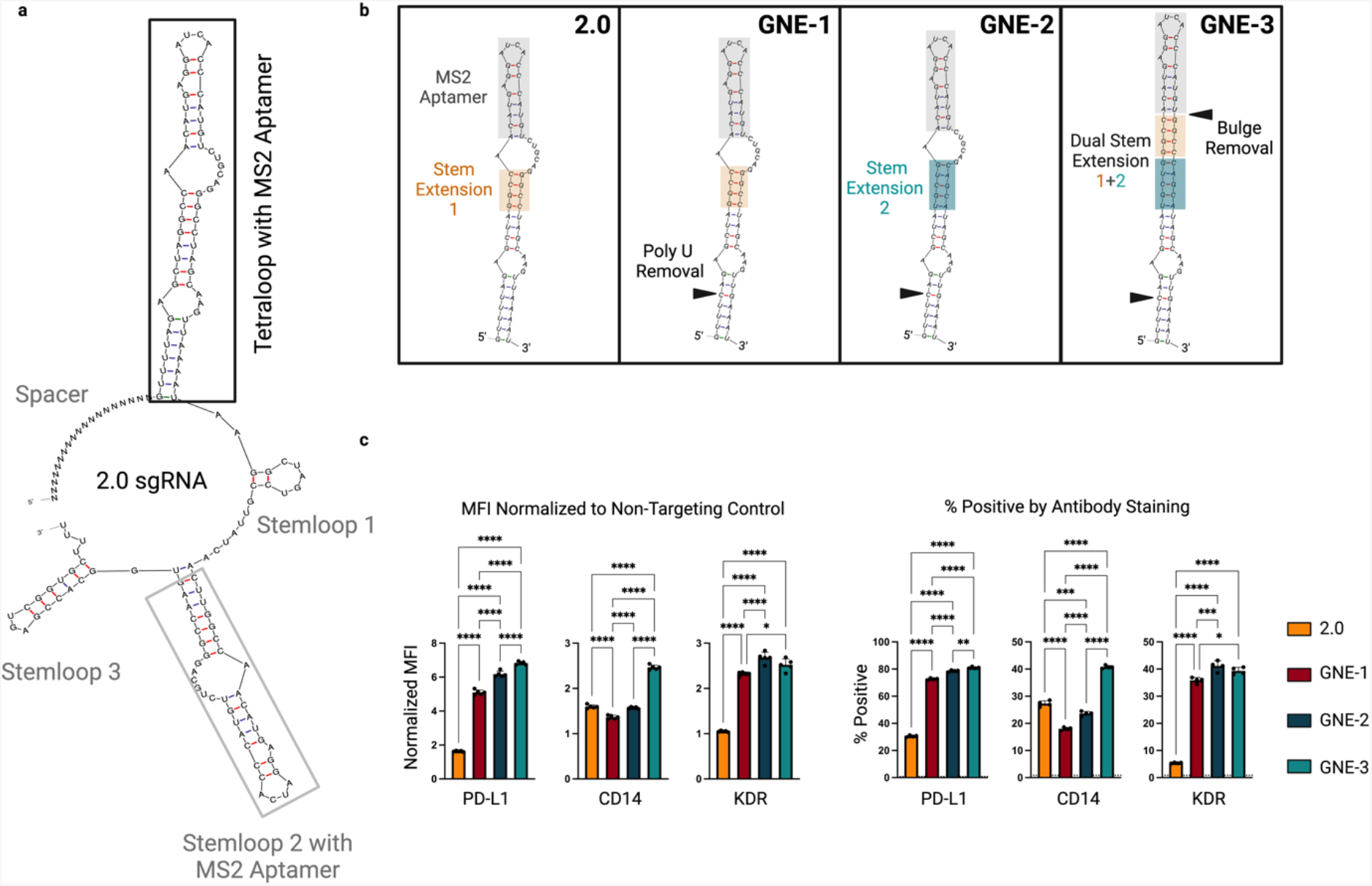
CRISPR activation efficiency of sgRNAs containing scaffold structural modifications. **a**, Structure of the 2.0 sgRNA^13^ with a modified MS2 aptamer containing tetraloop (black box) and stemloop 2 (gray box). **b**, Enlargement of tetraloop structure with highlighted sequence modifications in the alternate scaffolds evaluated. **c**, Flow cytometry data comparing activation efficiency of the four scaffold formats in a K562 CRISPRa-sel population. Cell populations were lentivirally transduced with the sgRNAs targeting 3 endogenous gene targets (PD-L1, CD14 or KDR) and analyzed 7 days post infection. Data represented as median fluorescence intensity (MFI) normalized to a cell population infected with a non-targeting sgRNA (left) or percentage positive (right) with non-targeting gRNA represented by dashed horizontal line. n=4 technical replicates per condition. Statistical comparison was performed by an unpaired 1-way ANOVA. * p<0.5, ** p<0.01, *** p<0.001. KDR-Kinase Insert Domain Receptor (VEGF2R/FLK1).

**Extended Data Fig. 4:**
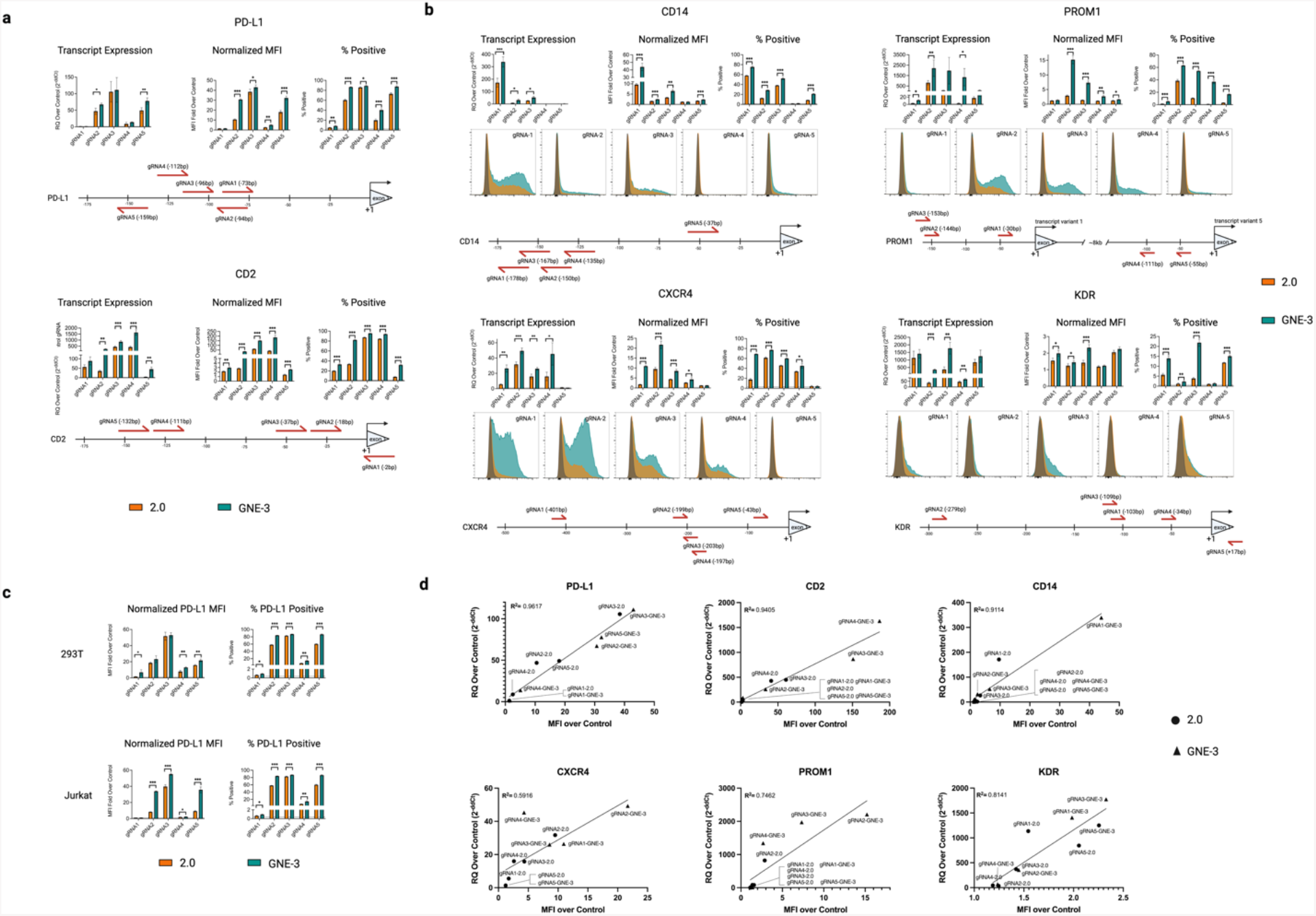
sgRNA activation efficiency of guides in the 2.0 or GNE-3 scaffold context. **a**,**b**, CRISPRa target gene activation by sgRNAs in a 2.0 (orange) or GNE-3 (teal) sequence context in a K562 CRISPRa-sel population. **(a)** Expression of PD-L1 (upper) or CD2 (lower) target genes assessed by qRT-PCR (left bar plots) and relative to a non-targeting control. Target gene expression assessed by flow cytometry and displayed by median fluorescence intensity (MFI) normalized to a non-targeting control (middle bar plot) or percent target positive (right bar plot). Background percent positive using a non-targeting control sgRNA indicated by horizontal dashed line. The position of guide RNA binding relative to the transcription start site (TSS) for each gene is indicated below. **(b)** Activation of 4 additional target genes with 2.0 or GNE-3 sgRNAs as assessed by transcript expression (top left), or flow cytometry (Normalized MFI-center; percent positive right or representative histograms, bottom panels). Guide position for each gene relative to the TSS indicated below. **c**, CRISPR mediated activation of PD-L1 in two additional cell lines (293T-top and Jurkat-bottom) by sgRNAs in a 2.0 or GNE-3 sequence context. PD-L1 expression assessed by flow cytometry and represented as MFI normalized to a non-targeting control (left) or percentage PD-L1 positive (right). Percent PD-L1 positive of cells infected with a non-targeting sgRNA represented by a horizontal dashed line. **d**, Scatter plots showing correlation of protein expression (normalized MFI) and transcript expression (qRT-PCR) for each sgRNA evaluated. R squared for simple linear regression analysis indicated. Statistical significance determined by a 2-tailed Student’s t-test assuming unequal variance. * p<0.5, ** p<0.01, *** p<0.001. RQ-Relative Quantity

**Extended Data Fig. 5:**
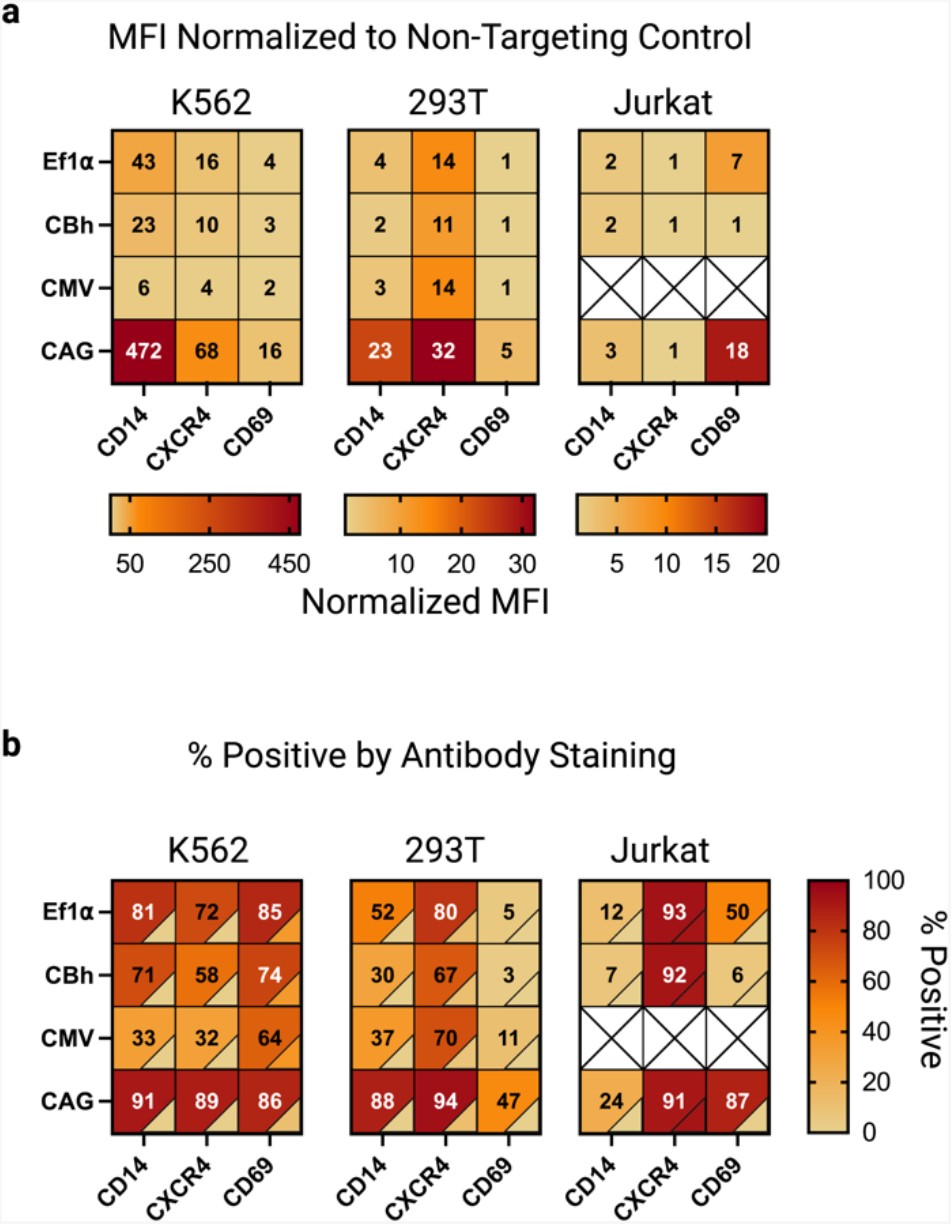
Extended CRISPRa-sel promoter optimization and application in K562, 293T and Jurkat cell lines. **a**,**b** Heatmap representing endogenous gene activation of 3 target genes in K562, 293T or Jurkat populations engineered with CRISPRa-sel piggyBac vectors driven by the indicated promoters (Ef1a, CBh, CMV or CAG). Cells were infected with lentivirus encoding dual GNE-3 sgRNAs targeting the indicated genes (CD14, CXCR4 or CD69) and assayed by flow cytometry 14 days post-infection/zeo selection. (**a**) Median fluorescence intensity (MFI) normalized to a stained cell population infected with a non-targeting control sgRNA in K562 (left), 293T (center) or Jurkat (right). Colorometric scale for each cell line indicated. (**b)** Percentage of the indicated cell populations positive by antibody staining. Percent positive of stained cell populations expressing a non-targeting sgRNA represented colorimetrically in the lower right corner of each cell. CRISPRa cell populations generated in triplicate and infected with indicated sgRNAs in technical duplicates.

**Extended Data Fig. 6:**
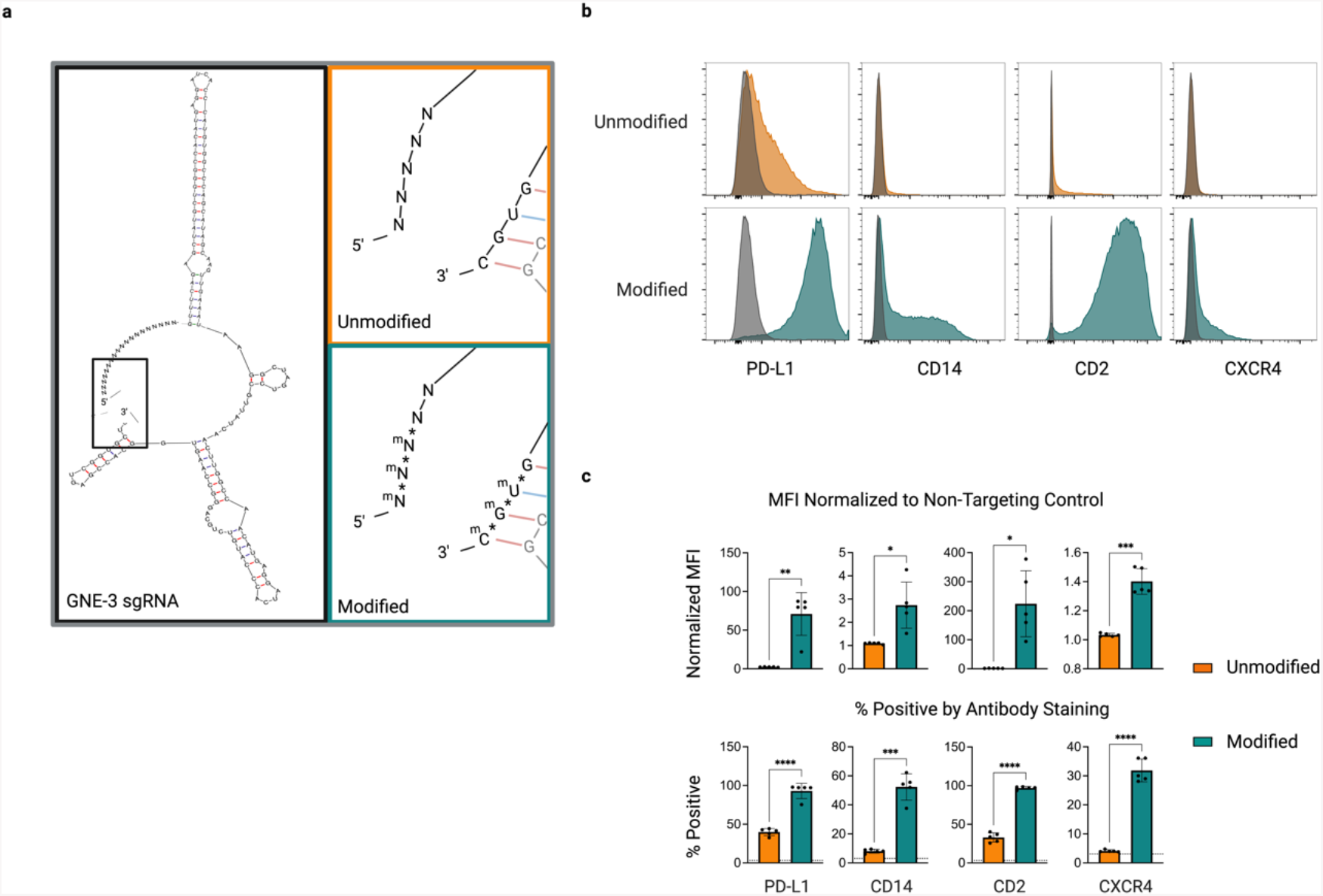
Chemical modification of synthetic GNE-3 sgRNAs enhances target gene activation. **a**, Structural diagram of a full sgRNA with GNE-3 scaffold highlighting 5’/3’ ends (black box). Magnified view of sgRNA 5’/3’ end regions highlighting unmodified (orange) or modified (teal) nucleotides. Modified sgRNAs contain 2’-O-methyl (m)/ phosphorothioate (*) linker modifications. **b**,**c** Assessment of CRISPR mediated gene activation by unmodified or modified sgRNAs in a CAG-CRISPRa-sel engineered K562 population and assessed 3 days post-sgRNA delivery. (**b)** Gene expression displayed by representative flow cytometry histograms in populations electroporated with unmodified (top row, orange) or modified (bottom row, teal) GNE-3 sgRNAs. Stained cells electroporated with a non-targeting synthetic sgRNA overlaid in gray. **(c)** Median fluorescence intensity (MFI) of K562 populations stained with antibodies for the indicated genes (PD-L1, CD14, CD2, CXCR4) and normalized to a population of stained cells electroporated with a non-targeting sgRNA (top). Percentage of cells positive by antibody staining (bottom). Background staining of a cell population electroporated with a non-targeting control sgRNA indicated with a dashed horizontal line. n=5 technical replicates per condition. Statistical significance determined by an unpaired 2-tailed t-test with a Welch’s correction. * p<0.5, ** p<0.01, *** p<0.001.

**Extended Data Fig. 7:**
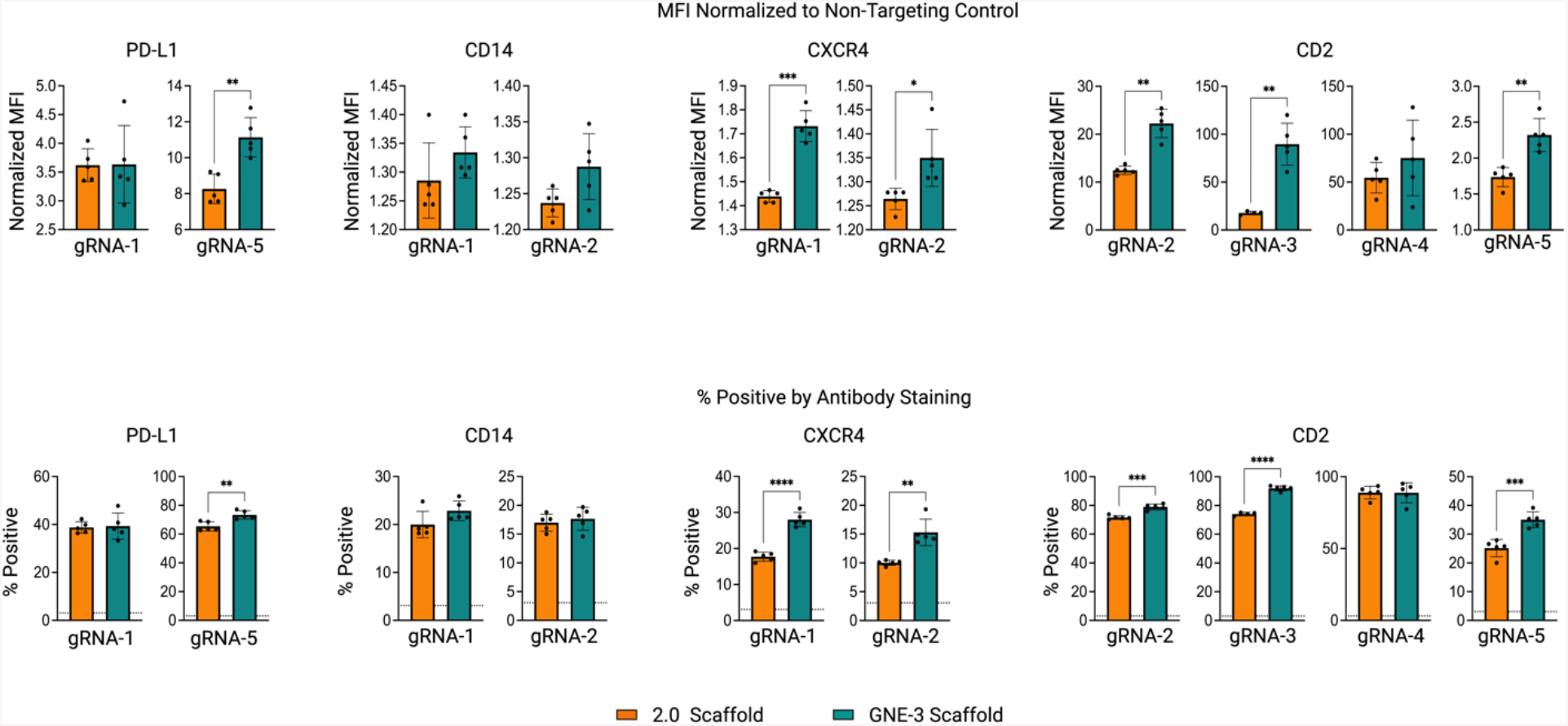
Activation efficiency of synthetic, modified, sgRNAs with a 2.0 or GNE-3 scaffold. Evaluation of CRISPR mediated gene activation in a CAG-CRISPRa sel engineered K562 population electroporated with modified synthetic sgRNAs in a 2.0 (orange) or GNE-3 (teal) scaffold context. Activation of 4 target genes (PD-L1, CD14, CXCR4 or CD2) by flow cytometry 3 days post sgRNA electroporation. Data represented as normalized median fluorescent intensity (top) or percent positive observed by antibody staining (bottom). Background antibody staining indicated by horizontal dashed line. n=5 technical replicates per condition. Statistical significance determined by an unpaired 2-tailed, t-test with a Welch’s correction. * p<0.5, ** p<0.01, *** p<0.001.

**Extended Data Fig. 8:**
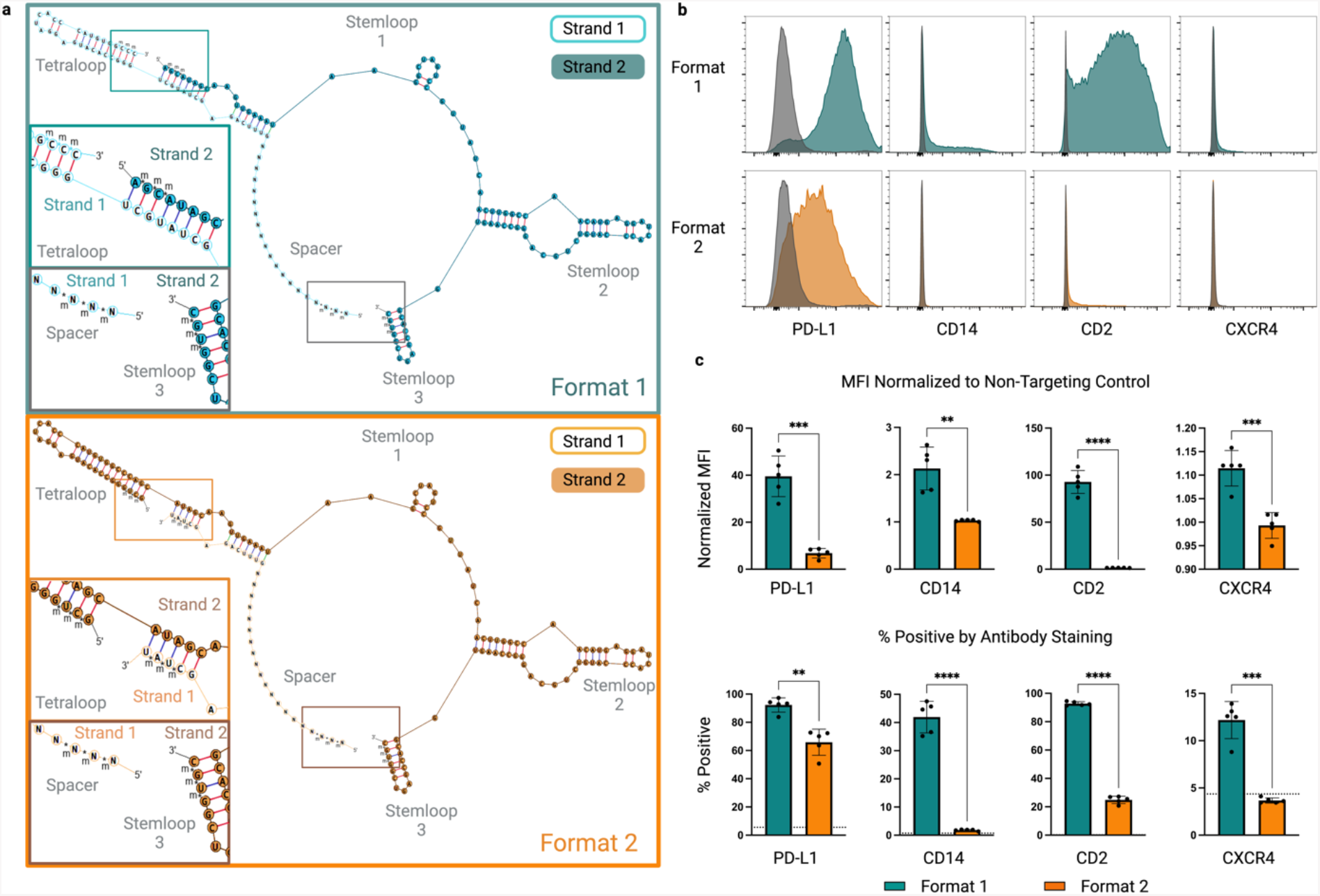
Comparison of alternate synthetic, 2-part GNE-3 guide RNA formats. **a**, Diagrams of alternate 2-part, modified, GNE-3 scaffold containing guide RNA structures. Format 1 (teal) encodes a spacer and the majority of the MS2 containing tetraloop on strand 1. Strand 2 of this format encodes the remainder of the tetraloop as well as stemloop 1, stemloop 2 with a second MS2 aptamer and stemloop 3. Strand 1 of the alternate format 2 structure (orange) encodes a spacer sequence and a short 3’ sequence predicted to anneal to the strand 2 scaffold containing the GNE-3 tetraloop and stemloops 1-3. Inset panels indicate the 2’-O-methyl and phosphorothioate linkers on the distal ends of each strand. **b-c**, Evaluation of alternate formats in CAG CRISPRa-sel engineered K562 populations and 3 days post gRNA electroporation. Expression of indicated endogenous target genes (PD-L1, CD14, CD2 or CXCR4) were evaluated by flow cytometry and presented as **b**, representative histograms overlaid with non-targeting control electroporated populations (gray) or **c**, summarized as normalized median fluorescence intensity (top) or percentage target gene positive (bottom). Background staining indicated by horizontal dashed line. n=5 technical replicates per condition. Statistical significance determined by an unpaired 2-tailed t-test with a Welch’s correction. * p<0.5, ** p<0.01, *** p<0.001. m=2’-O methyl. *= phosphorothioate linker.

**Extended Data Fig. 9:**
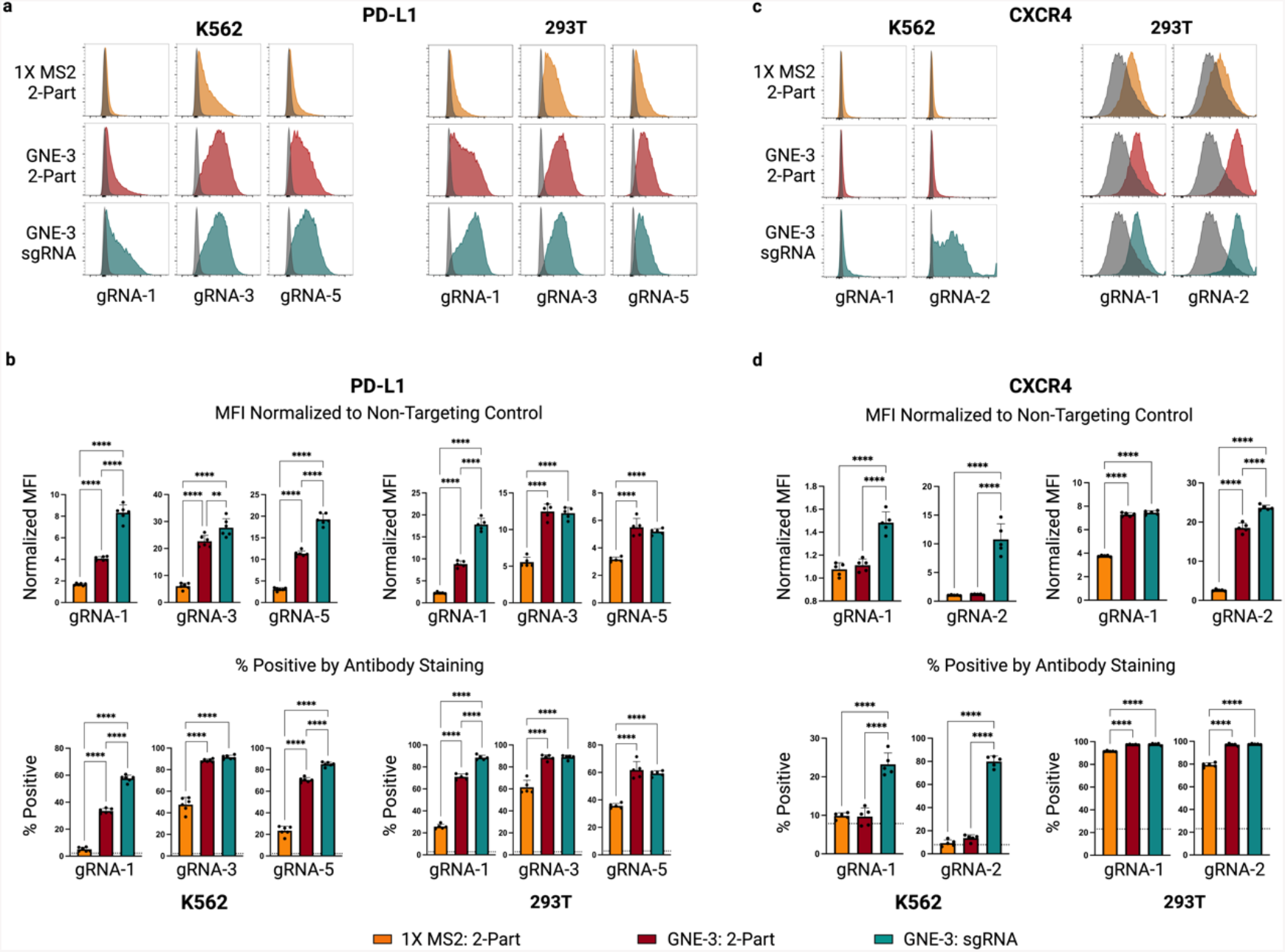
Evaluation of CRISPRa synthetic guide formats in an expanded cell panel. **a-d**, Target gene activation comparing three different synthetic guide RNA formats, 1X MS2 2-Part (orange), GNE-3 2-Part (maroon) and GNE-3 sgRNA (teal) as illustrated in Fig. 4a, across CRISPRa-sel K562 and 293T cell populations. **(a)** PD-L1 and **(c)** CXCR4 activation assessed by flow cytometry of antibody-stained cell populations 3 days post gRNA electroporation (K562) or transfection (293T). Representative histograms for each gene targeting gRNA are shown, overlaid with histograms for the non-targeting gRNA control (grey). Median fluorescence intensity for each gene targeting gRNA normalized to the non-targeting gRNA control for (**b**, top) PDL1 and (**d**, top) CXCR4. Percentage of cells positive by antibody staining for PD-L1 (**b**, bottom) or CXCR4 (**d**, bottom), with background staining indicated by the horizontal dashed line. n=5 technical replicates per condition. Statistical comparison was performed by an unpaired 1-way ANOVA. ** p<0.01, **** p<0.0001.

